# Two case studies detailing Bayesian parameter inference for dynamic energy budget models

**DOI:** 10.1101/259705

**Authors:** Philipp H. Boersch-Supan, Leah R. Johnson

## Abstract

Mechanistic representations of individual life-history trajectories are powerful tools for the prediction of organismal growth, reproduction and survival under novel environmental conditions. Dynamic energy budget (DEB) theory provides compact models to describe the acquisition and allocation of energy by organisms over their full life cycle. However, estimating DEB model parameters, and their associated uncertainties and covariances, is not trivial. Bayesian inference provides a coherent way to estimate parameter uncertainty, and propagate it through the model, while also making use of prior information to constrain the parameter space. We outline a Bayesian inference approach for energy budget models and provide two case studies – based on a simplified DEBkiss model, and the standard DEB model – detailing the implementation of such inference procedures using the open-source software package deBInfer. We demonstrate how DEB and DEBkiss parameters can be estimated in a Bayesian framework, but our results also highlight the difficulty of identifying DEB model parameters which serves as a reminder that fitting these models requires statistical caution.

## 1 Introduction

Dynamic energy budget (DEB) theory (Kooijman, 2010) provides a powerful and well tested framework to describe the acquisition and use of energy by individual organisms over their entire life cycle. The standard DEB model is built on rules inherent to the process of resource uptake and allocation by all heterotrophs. It is a compact model that is able to describe the full life cycle bioenergetics of a living animal (Kooijman, 2010). DEB models are used as tools to address both fundamental and applied questions in ecology, e.g. about metabolic scaling (Maino et al., 2014), life-history strategies (Kooijman, 2013), in ecotoxicology (Billoir et al., 2008b; Jager et al., 2006; Jager and Zimmer, 2012), or as components of population models (Billoir et al., 2007; Martin et al., 2012; Smallegange et al., 2017).

Because of strong correlations between individual parameters, simultaneous parameter inference for DEB models is not trivial (Billoir et al., 2008a; Johnson et al., 2013). The difficulty of estimation is by no means unique to DEB models, but is commonly encountered in dynamic systems models across scientific disciplines (Aster et al., 2011; Brewer et al., 2008; Johnson and Briggs, 2011). Parameter inference for DEB model parameters themselves is often based on a non-linear least squares regression approach, the so-called covariation method (Lika et al., 2011). The covariation method makes use of constraints on parameters that follow from theory (Lika et al., 2014, 2011), as well as by reducing the effective number of parameters by the use of so-called pseudo data: canonical values for certain parameters which enter the estimation procedure with low weights. This approach has been successfully used to parameterize DEB models for over 1000 species (Marques et al., 2018). However, one drawback of the method is that uncertainty estimates of parameters are not readily available. Furthermore, while the DEB literature acknowledges the importance of biological variability (e.g. Lika et al., 2014), input data are treated as known without error for the purposes of the parameter estimation. While measurement error for many observable traits used to parameterize DEB models may indeed be small, trait data often exhibits heterogeneity between individuals of a species, which would be expected to reflect individual heterogeneity in the underlying metabolic parameters. Given the potential of DEB theory as a building block for population models, we feel these are important hurdles to overcome, so more value can be added to DEB-based population models by incorporating both better estimates of parameter uncertainty, and a better reflection of individual variability.

In contexts where quantification of uncertainty in parameters is desired, the Bayesian inference framework offers multiple advantages. First, multiple sources of uncertainty can be accounted for. Second, the use of informative priors can constrain the parameter space to biologically sensible outcomes (e.g. by constraining maximum lengths or reproductive rates to values that are realistically attainable by a given species), while allowing fairly straightforward assessment of the influence of the prior information. Finally, the implementation of hierarchical models which allow inferences about both individual and population heterogeneity, as well as partial information pooling across individuals, is conceptually straightforward (Gelman et al., 1996). Computationally, hierarchical inference in differential equation models still provides a number of challenges (see e.g., Krauss and Schuppert, 2017). It is therefore beyond the scope of this tutorial.

Bayesian parameter inference for DEB models has been demonstrated by Billoir et al. (2008b) and Johnson et al. (2013). However, until recently, general inference for these models in a Bayesian framework has required that the practitioner be able to implement the full inference procedure, from the differential equation model specification through to the sampler used to obtain posterior draws. Here, we present a tutorial for the estimation of model parameters for dynamic energy budget models using the open-source R package deBInfer (Boersch-Supan et al., 2017) which makes the approach simpler to implement.

We present a tutorial detailing Bayesian parameter inference for two case studies. The first is based on a DEBkiss model, a simplified bioenergetic model that builds on DEB theory (Jager et al., 2013; Jager, 2016). We follow this with a case study based on the standard DEB model. In each case study we discuss how the model is implemented and the approach estimate parameters. The R and C code needed to reproduce all of the analyses are available as supplementary materials.

Given the complexity of both DEB theory itself, and the statistical and computational procedures involved in parameter inference for differential equations, this tutorial presupposes a basic knowledge of DEB theory, and a basic familiarity with the R computing environment. For background materials on DEB theory we refer the reader to Maino et al. (2014) as an initial introduction, and the recent review by Jusup et al. (2017) and the monograph by Kooijman (2010) for a comprehensive treatment. General approaches to working with differential equation models in R are outlined in Soetaert et al. (2010), and comprehensively in the monographs Soetaert and Herman (2008); Soetaert et al. (2012).

## 2 Basics of the Bayesian Approach

Bayesian approaches for parameter estimation in complex, nonlinear models are being applied with increasing frequency to a broad range of biological models (e.g. Coelho et al., 2011; Voyles et al., 2012; Johnson et al., 2013; Smith et al., 2015). Here we very briefly explain the rationale behind the approach. Further details on the statistical background and software implementation can be found in the literature, (e.g. Clark, 2007; Gelman et al., 2003; Hobbs and Hooten, 2015; Johnson et al., 2013; Boersch-Supan et al., 2017).

In the Bayesian approach the model, and in particular its parameters, are viewed as random variables. In contrast, the data, once observed, are treated as fixed. This approach to parameter inference is attractive, as it provides a coherent framework that allows the incorporation of uncertainty in the observation process and model parameters, and it relaxes the assumption of normal errors that is inherent in least-squares fitting. It provides us not only with full posterior probability distributions describing the parameters, but also with posterior distributions for any quantity derived from the parameters, including the model trajectories. Further, the Bayesian framework naturally lets us incorporate prior information about the parameter values and examine the sensitivity of our inferences to this incorporated information. This is particularly useful in the context of DEB theory, where there are known biological and theoretical constraints on parameters (Lika et al., 2011, 2014; Johnson et al., 2013). For example, many biological quantities, such as body size, are strictly non-negative, and most DEB parameters have at least one well defined theoretical bound, e.g. the allocation fraction *κ* is bounded on the interval [0,1]. Using informative priors can help constrain the parameter space, aiding parameter identifiability.

The purpose of our case studies is to show how to estimate the joint posterior probability distribution of the parameters of an energy budget model 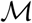, given an empirical dataset 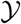, and accounting for the uncertainty in the data. The deterministic model 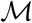 takes the general form

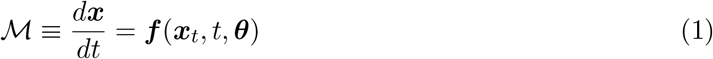

where ***x*** is a vector of state variables evolving with time; ***f*** is a functional operator that takes a time input and a vector of continuous functions ***x**_t_*(*θ*) and generates the vector 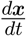 as output; and ***θ*** denotes a set of parameters.

Using Bayes’ Theorem (Hobbs and Hooten, 2015) we can calculate the posterior distribution of the model parameters, given the data and the prior information as

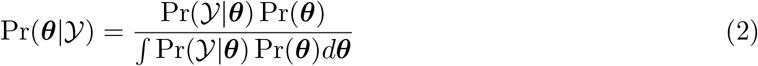

where Pr() denotes a probability, 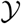 denotes the data, and ***θ*** denotes the set of model parameters. The product in the numerator is the *joint distribution,* which is made up of the *likelihood* 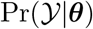 (corresponding to the *likelihood function* 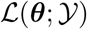 of the frequentist approach), which gives the probability of observing 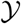 given the deterministic model 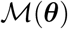, and the *prior distribution* Pr(***θ***), which represents the knowledge about ***θ*** before the data were collected.

Closed form solutions for the posterior are practically impossible to obtain for complex nonlinear models such as DEB models, but they can be approximated numerically, e.g. by using Markov Chain Monte Carlo (MCMC) integration methods (Gilks et al., 1995).

### 2.1 Computation using the deBInfer package

The deBInfer package (Boersch-Supan et al., 2017) provides templates for implementing dynamical models consisting of a deterministic differential equation (DE) model and a stochastic observation model and subsequently for performing inference for these models. To perform inference, R functions or data structures must be specified to represent both the dynamical (here bioenergetic) model and the observation model (i.e., the data likelihood). Further, all model and observation parameters must be declared, including prior distributions for those parameters that are to be estimated or values for those that are fixed. The DE model itself can also be provided as a shared object, e.g. a compiled C function, which can considerably speed up inference (see Appendix S3 in Boersch-Supan et al., 2017). deBInfer then samples from the posterior distributions of parameters via MCMC, specifically using Metropolis-Hastings updates nested within a Gibbs sampler (Brooks et al., 2011). As each sample of the posterior distribution is obtained, the differential equation model must be solved numerically within the MCMC procedure.

## 3 Case Study 1: DEBKiss Model

The standard DEB model is a powerful framework to describe the bioenergetics of an organism across its full life cycle (Kooijman, 2010). However, that power comes at a cost of many complex equations with many parameters needing a great deal of data to parameterize properly. In an effort to develop a simpler model that still exhibits important features of the full DEB theory Jager et al. (2013) developed the DEBKiss model. It is a model inspired by DEB but “with a strong focus on the KISS principle” (Jager et al., 2013). The main departures from DEB are the absence of a reserve buffer and a maturation state variable. This has the effect of reducing the number of differential equations in the system, as well as reducing the number of parameters. The model is slightly less flexible. For instance, the organism must mature at a fixed length or weight. In contrast, the DEB framework allows maturation to happen once sufficient complexity has been accrued, which can correspond to different weights or lengths in organisms living in differing food environments.

We use the DEBKiss model as a simplified DEB-like model to show the basics of the Bayesian approach to fitting models of this sort. We perform inference using a subset of the data set used in the paper introducing DEBKiss: data on growth (measured as shell length in mm) and reproduction (cumulative egg production) of the pond snail, *Lymnaea stagnalis*. These data come from a series of partial life experiments. Juvenile snails that were the same age (113 days from when the egg was laid) and approximately the same size were selected and followed for an additional 140 days (data from Zimmer et al., 2012). To keep this example especially simple we use data from a single food level treatment, specifically snails that were fed *ad libitum* over the course of the experiment. Further, we only estimate a subset of the parameters estimated in the original DEBKiss paper (which were, in turn, a subset of all of the parameters) as we found during initial analyses that not all parameters were practically identifiable with data for a single food level under the assumption of unknown observation error. The model is specified and described in detail in both the main and supplementary text of Jager et al. (2013), thus we do not provide a complete overview here, instead summarizing a few key equations and parameters. The DEBkiss model is formulated as a set of coupled ordinary differential equations for 3 state variables: the egg buffer *W_B_*; structural body mass *W_V_*; and reproduction buffer *W_R_*. The structural body mass is related to the physical length, 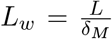 where 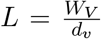, and all equations may be written in terms of these lengths instead of the masses. Thus the equations governing the growth and and allocation to reproduction can be written as

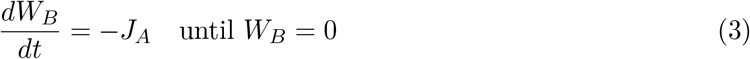

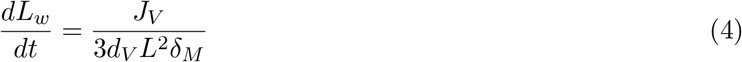

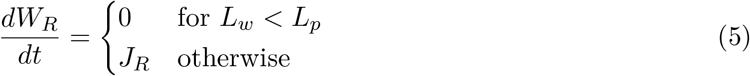

with parameters defined in Table 1. The data we utilize is recorded beginning after hatching, and thus we begin with the reserve in the egg, *W_B_* = 0. The complete implementation of the model in R, including the inference shown here, is included as supplementary materials.

### 3.1 Bayesian Parameter Estimation

For simplicity, we focus on estimating a subset of parameters from data on snail growth and reproduction at a single food level, keeping most model parameters fixed (see Table 1). The DEBKiss model was implemented as a set of differential equations following Jager et al. (2013). Similarly to the standard DEB model, the state variables in the DEBKiss framework are not all directly measurable. However, it is possible to specify how measured quantities, such as length and numbers of eggs, are related to the state variables. For this application, we used the formulation of the DEBKiss equations in terms of physical length and cumulative number of eggs produced by the snails.

#### Likelihood

Once the differential equations have been specified, the likelihood of the data conditional on the parameters and model must next be specified. The `deBInfer` package allows substantial flexibility in the probability distributions that may be used for this purpose. For instance, one could allow normal errors for lengths and truncated normal or log-normal, or Poisson errors for egg counts. This allows the user to choose an appropriate distribution that is consistent with the characteristics of the data the user wishes to model. The snail data we use here consist of average lengths (mm) and mean cumulative eggs. Thus both the lengths and eggs are appropriately modeled as continuous distributions. Additionally both are constrained to be positive and have error that increases over time (as small differences between individuals is likely to be magnified as the grow).

To define our likelihood, we must relate our measured quantities to the model outputs. We assume that the observed length, 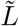, is the product of two quantities: the predicted physical length from the DEBKiss model, denoted as *L_w_* and a log-normally distributed, multiplicative noise term. Thus:

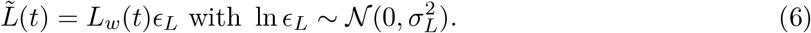

Similarly, the resources allocated to reproduction, *W_R_*, are related to the number of eggs. However, the number of eggs created depend on the energy needed per egg, *W_B0_*, and the conversion efficiency of assimilated energy to egg, *y_BA_*. Again, the noise is assumed to be multiplicative and log-normal (since we cannot use a Poisson or similar as the counts have been averaged), so the cumulative egg production at any given time, *F*, is given by

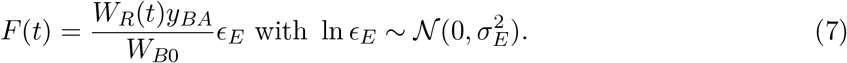

Conditional on the solution to the underlying differential equations we assume that the observed lengths and eggs are independent at each time. Thus the likelihood is constructed by multiplying the (independent) likelihoods of the lengths and fecundity at each time point together.

#### Priors and sampling details

We chose relatively un-informative priors for the four parameters that we chose to estimate. The choice of prior here was primarily guided by simple constraints on the values that the parameters may take. For example, *κ*, the proportion of energy directed towards growth processes, must lie between 0 and 1. Thus we used a uniform prior over this full range as the prior. Priors for estimated parameters are specified in Table 1.

**Table 1:** State variables and parameters for the DEBKiss model, as well as fixed values and prior distributions used for the parameter inference procedure.

| Parameter | Description | Fixed value or Prior distribution | Reasoning |
| --- | --- | --- | --- |
| $W_B$ | Mass of egg buffer | | |
| $W_{B0}$ | Mass of assimilates in an egg | 0.15 mg | |
| $F$ | Cumulative egg production | | |
| $L_w$ | Physical body length | | |
| $W_R$ | Mass of reproductive buffer | | |
| $J_A$ | Mass flux for assimilation $fJ_{Am}^a L^2$ | | |
| $J_V$ | Mass flux for structure $y_{VA}(\kappa J_A - J_M)$ | | |
| $J_M$ | Mass flux for somatic maintenance $J_M^v L^3$ | | |
| $J_R$ | Mass flux to reproductive buffer $(1 - \kappa)J_A$ | | |
| $J_{Am}^a$ | Surface-specific maximum assimilation rate | 0.11 mg/mm<sup>2</sup>/d | |
| $d_V$ | Dry-weight density of structure | 0.1 mg/mm<sup>3</sup> | |
| $L$ | Volumetric body length | | |
| $\delta_M$ | Shape correction coefficient | 0.40 | |
| $f$ | Functional response | 1 | |
| $y_{VA}$ | Yield of structure on assimilates | 0.8 | |
| $y_{BA}$ | Yield of egg buffer on assimilates | 0.95 | |
| $\kappa$ | Fraction of assimilation flux for soma | Uniform<br>$a = 0; b = 0$ | bounded on [0,1] |
| $\ln(J_M^v)$ | ln of volume-specific maintenance costs | normal<br>$\mu = 0; \sigma^2 = 100$ | weakly informative prior, constraining the untransformed parameter to be positive. |
| $\ln \epsilon_L$ | ln of observation error on length | lognormal<br>$\mu = 0; \sigma^2 = 1$ | weakly informative prior regularizing towards 0 |
| $\ln \epsilon_E$ | ln of observation error on F | lognormal<br>$\mu = 0; \sigma^2 = 1$ | weakly informative prior regularizing towards 0 |

In addition to a prior distribution, the user must specify a *proposal* distribution for each parameter being sampled (Gilks et al., 1995). This distribution determines how new values of each parameter are chosen as the MCMC algorithm progresses. In the `deBInfer` package one can choose between 3 options: 1) a random walk proposal with a normal proposal distribution 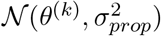, centered at the current accepted parameter value *θ^(k)^*; 2) a random walk proposal with a uniform distribution 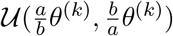 that is asymmetric around the current accepted value *θ^(k)^*; 3) independent draws from the prior distribution. In the example here we chose all random walk proposals. For *k* and 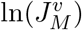 we used a normal proposal. The normal proposal distribution is generally a robust default choice. However, more efficient sampling can be achieved in certain situations, in particular where a parameter is strictly positive. This is why we used asymmetric uniform proposals for the two standard deviations, which by definition have a lower bound at zero. The asymmetric uniform proposal is especially good for sampling parameters that are bounded on the left with values that may be close to that lower bound as it ensures strictly positive proposals and smaller sampling increments towards the bound.

### 3.2 DEBKiss Model: Results

#### MCMC Output Diagnostics

When examining the posterior output from the MCMC produced by `deBInfer`, the first step is always to check for *mixing* and *convergence* of the MCMC chain by plotting traces of the chains (e.g., Figure 1). A “good”, well behaved chain (i.e., that mixes adequately and that has converged to the posterior distribution) is sometimes described as resembling a “fuzzy caterpillar”. A chain that doesn’t look very fuzzy is often called a “sticky” chain – it has high auto-correlation and the effective sample size is low. If the chain has converged a horizontal line should approximately go through the center of the trace and there shouldn’t be major patterns, such as a linear trend, visible. The chains for this example appear to be well behaved, and seem to indicate both adequate mixing and convergence. The subtleties of assessing mixing and convergence is beyond the scope of this paper, but may be found in textbooks such as Gilks et al. (1995) or Hobbs and Hooten (2015). A more formal assessment of convergence can be obtained by running multiple MCMC chains and calculating the potential scale reduction factor, a measure comparing within-chain and between-chain variances (Brooks and Gelman, 1998). Approximate convergence is diagnosed when the upper limit of this measure is close to 1 for each variable.

**Figure 1:**
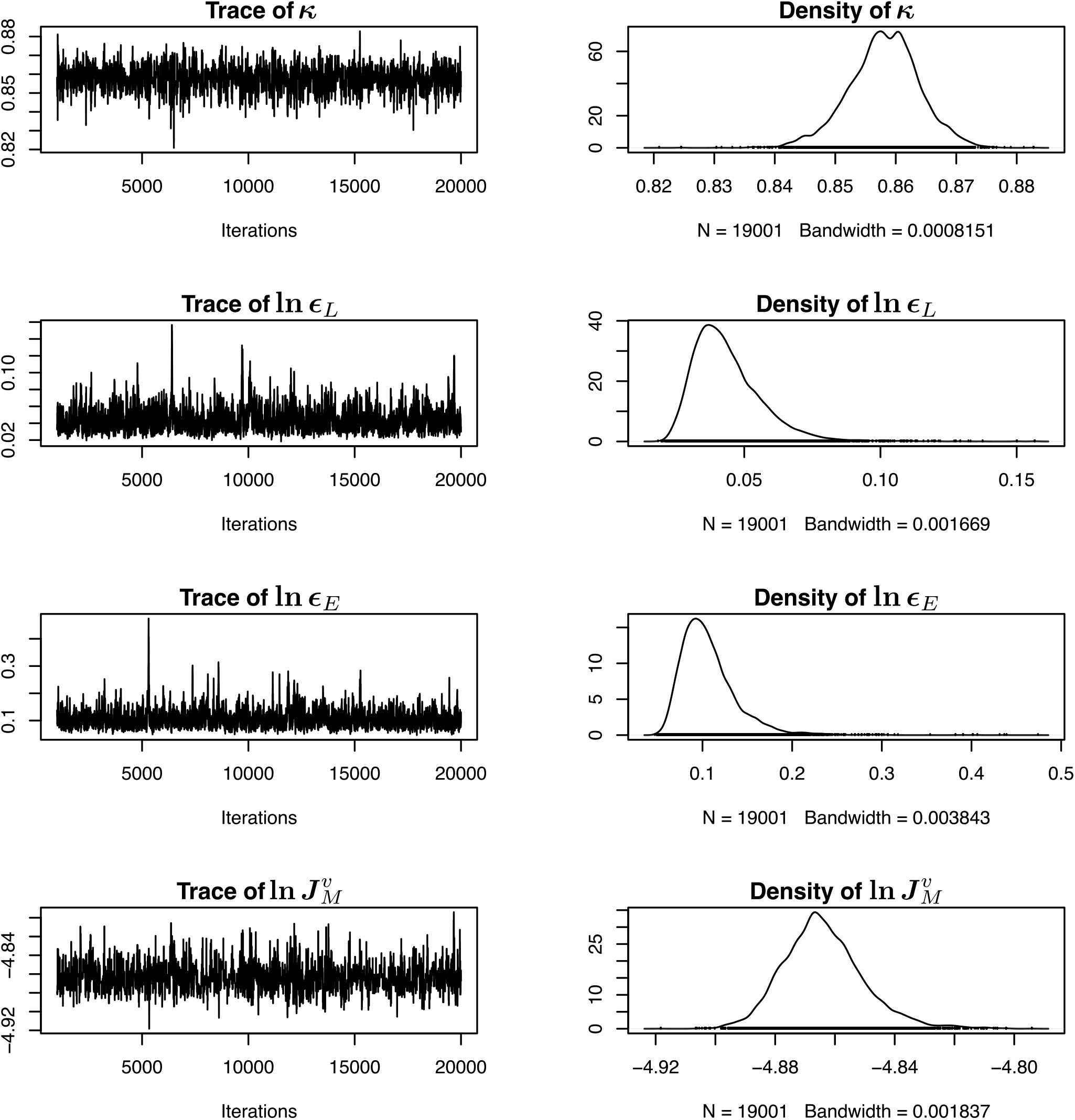
The MCMC traces and marginal distribution for two estimated observation parameters (the noise terms on the observed length measurements and egg counts) and two estimated parameters of the deterministic DEBKiss model indicate convergence and good mixing.

Once mixing and convergence have been assessed, the next, very important, diagnostic to check is the prior-posterior overlap. Priors in Bayesian analyses can be double edged swords – they allow us to incorporate previous knowledge and constraints into our process in a formal way. However it is possible to inject more prior information than one means to through the prior. If you don’t have good information about a parameter value, you ideally want to choose a “vague” prior so that the information in your data can drive the posterior results. Thus it is always a good idea to plot the marginal posterior distribution together with the marginal prior to confirm that your choice of prior is not influencing your posterior more than you intended. In our example, even though we knew the values that Jager et al. (2013) had previously reported for both DEBkiss parameters *k* and 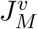, we wanted to incorporate as little additional information as possible in our priors. In Figure 2 we show the marginal posterior distribution for each parameter (in black) overlaid with the prior distribution (in red). Notice that across all 4 panels the red line lies across the very bottom of the panel – the priors have very little mass in the areas corresponding to the posterior distribution. In all cases the data seem to be informative for the parameters and the posteriors different from the priors.

**Figure 2:**
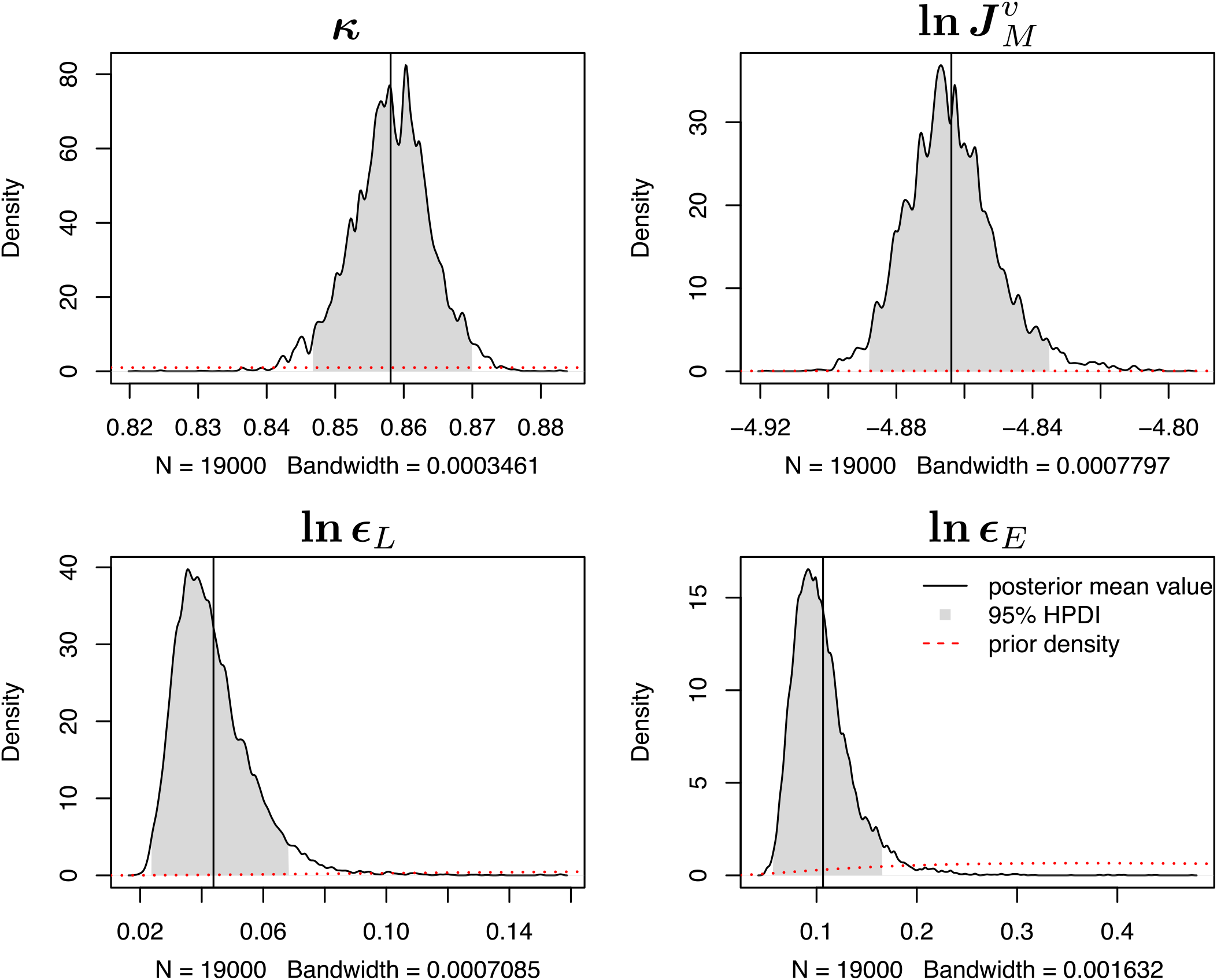
Marginal posterior samples of parameters (black lines) plotted with prior distributions (red). Shaded areas indicate the 95% highest posterior density (HPD) region. The posterior mean is indicated with a solid line. Notice that in all cases the prior is very different from the prior and the data are informative for all parameters.

We also typically examine the full joint posterior distribution by visualizing the pairwise joint distributions (e.g. Figure 3). The pairwise plots can give additional indications about the iden-tifiability of individual parameters, independent of the others. This is important, as in complex models not all parameters may be identifiable, that is different combinations of parameters lead to the same likelihood, making it impossible - with the data at hand - to decide among possible parameter values (Cobelli and Distefano, 1980). In this example we can see that the correlation between our estimated parameters is overall very low, with the strongest correlation being between *k* and 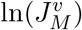, as we might expect as *κ* (the proportion of reserves invested in growth/maintenance) and 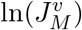 (volume-specific maintenance costs) together jointly determine the growth rate of the organism. For nonlinear systems such as these, often the observation parameters are not highly correlated with the primary parameters, but the model parameters themselves may be. Very strong correlations between parameters can indicate that the data are not fully informative for each parameter individually – for example it may be that the quotient or product is very tightly constrained by the available data, but the marginal uncertainty in the individual parameters is higher. This is not necessarily problematic, per se, but should be kept in mind when using and interpreting posterior samples, for example marginal posterior densities of individual parameters may be misleading about the possible values a parameter can take. Further, some patterns in the posterior, such as extreme nonlinear patterns (“banana” shapes, etc.) can indicate that parameters are strongly correlated and not well constrained. For an example of this for DEB models see Johnson et al. (2013).

**Figure 3:**
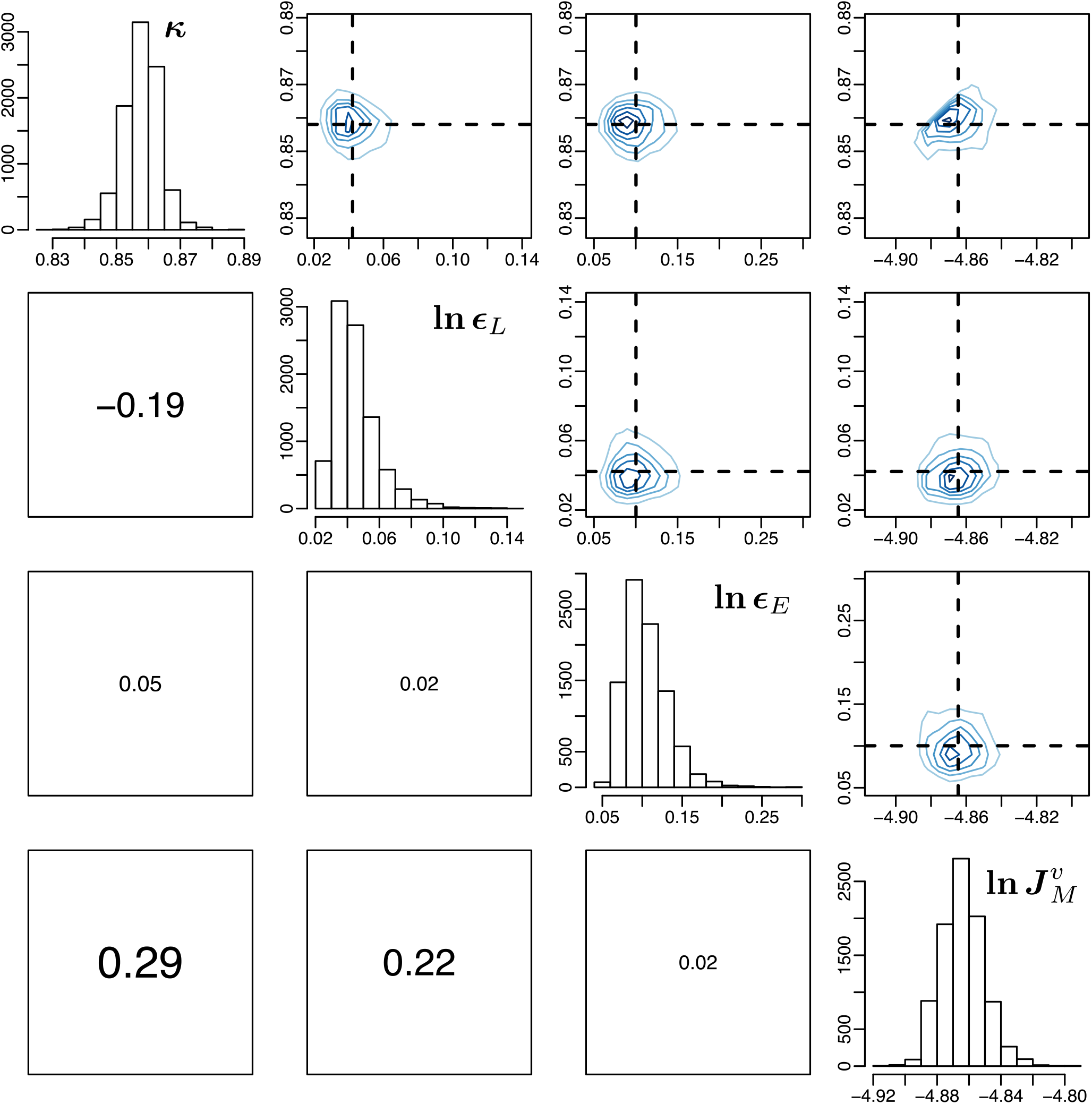
Joint samples from the full posterior of the 2 observation and 2 primary model parameters estimated for the DEBKiss model.

#### Posterior estimates of parameters

Now that we feel confident in the convergence of the chains and that our parameters are well identified we can interpret and present the inferred parameters, including the posterior uncertainties. Further, we can use the posterior distribution of parameters to obtain the posterior distributions of other functions of the parameters, such as the trajectories of the system.

In a Bayesian analysis we often report the marginal highest posterior density (HPD) interval or credible interval, which are the Bayesian analogs to confidence intervals. In Figure 2 we show a graphical representation of the HPD intervals for the parameters that we estimated. The shaded region corresponds to the HPD region (i.e., 95% of the posterior weight), and the HPD interval thus corresponds to the locations of the edges of the region. We indicate the posterior mean, often used as the point estimate for the parameter, using a solid line.

Finally, we can construct and visualize the posterior distributions of any functions of the parameters. For this example, we focus on the posterior distribution of the trajectories of the sets of differential equations. To obtain the posterior distribution of trajectories requires solving the set of differential equations with the parameters set to the estimated values in the posterior sample. For instance, in this example we collected *N* = 20000 samples of the posterior distribution of parameters. We discarded the first 1000 as burn-in (because for part of that the chain had not yet converged), leaving 19000 samples. We then “thinned” these samples (that is sub-sampled them), taking every 10th sample. This leaves 1900 parameter samples. For each of these samples we take the pair of primary parameter estimates together with the fixed parameters and initial conditions and solve the DEs. After repeating this for all 1900 samples we have 1900 trajectories of the system, reflecting parameter uncertainty of the deterministic model. We can obtain the mean behavior by taking the mean at each time point in the trajectory across the 1900 samples. Similarly we can calculate the credible intervals by obtaining the appropriate values of the quantiles at each time point. The posterior mean and credible intervals of the trajectories for our example are shown in Figure 4.

**Figure 4:**
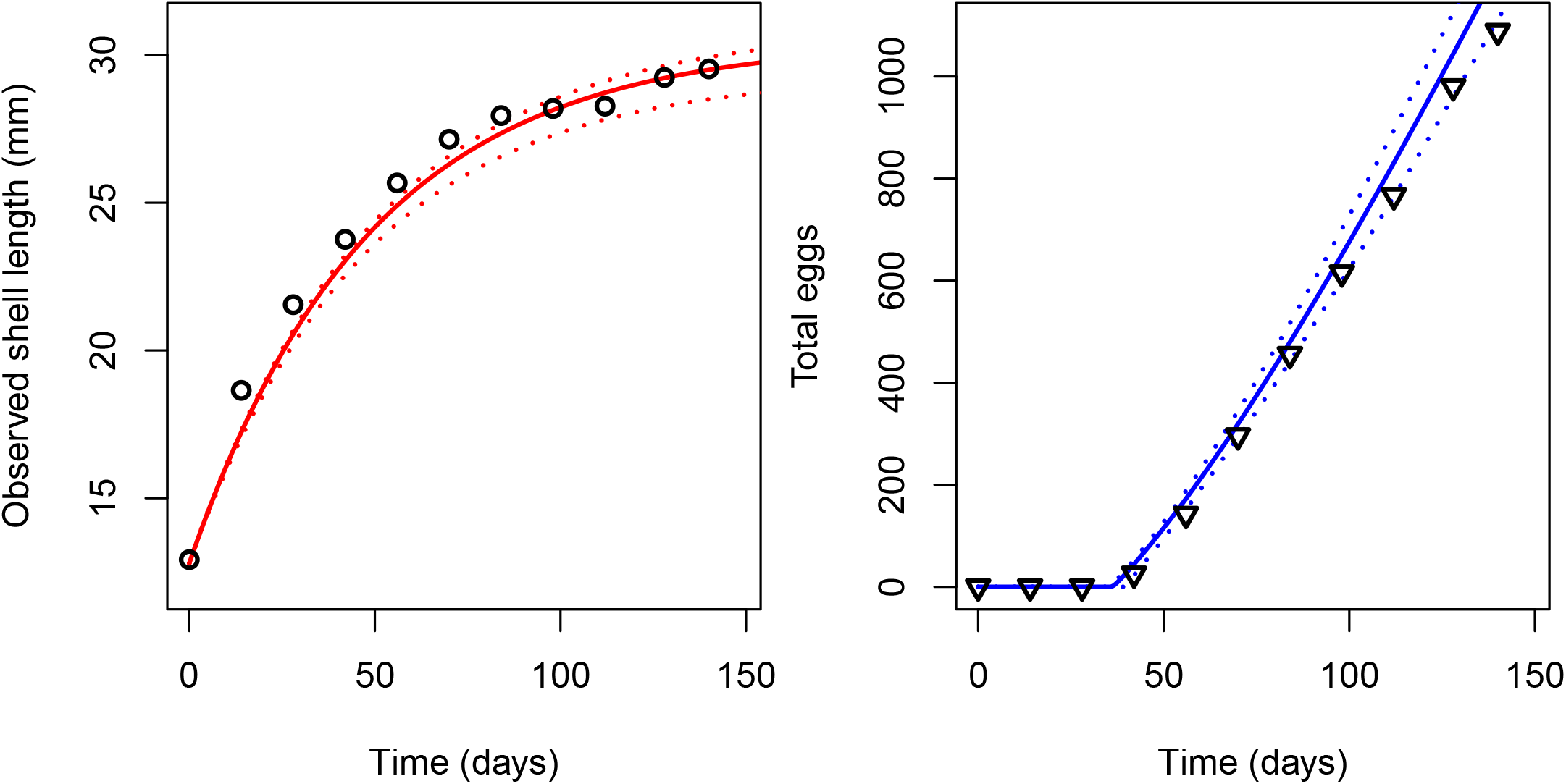
Snail growth (measured as shell length in mm) and reproduction data (from Zimmer et al., 2012) plotted with the posterior mean and 95% credible interval (CI) of the predicted shell length growth and cumulative egg production from the DEBKiss model. Solid lines indicate the posterior mean, and dotted lines the edges of the 95% CI.

Note that, unlike most methods for obtaining uncertainty estimates of parameters (e.g. obtained via maximum likelihood or least squares) we do not need to assume that the parameters are approximately multivariate normal. Although that assumption may not be far off for the simple example here, there are certainly cases where that assumption is a poor representation of the posterior distribution, and where assuming the parameters are jointly normal would result in very different predictions of the system trajectories and their uncertainties.

## 4 Case Study 2: The Standard DEB Model

For our second case study we attempt to replicate the DEBtool estimation of growth and reproduction parameters for the standard DEB model for the earthworm *Lumbricus terrestris*. We implemented the standard DEB model (Kooijman, 2010) in terms of scaled energy density e, scaled length *l*, and scaled maturity *u_H_* and reproductive buffer *u_R_*, respectively, as functions compliant with the ODE solvers provided by the deSolve package Soetaert et al. (2010). The model equations are given in Appendix A, parameter definitions are given in Tables 2 and 3. Further R functions to compute boundary values for the state variables from DEB parameters (Kooijman, 2009) were adapted from DEBtool routines and are available in the R package DEButilities which we provide in the supplementary materials.

**Table 2:**
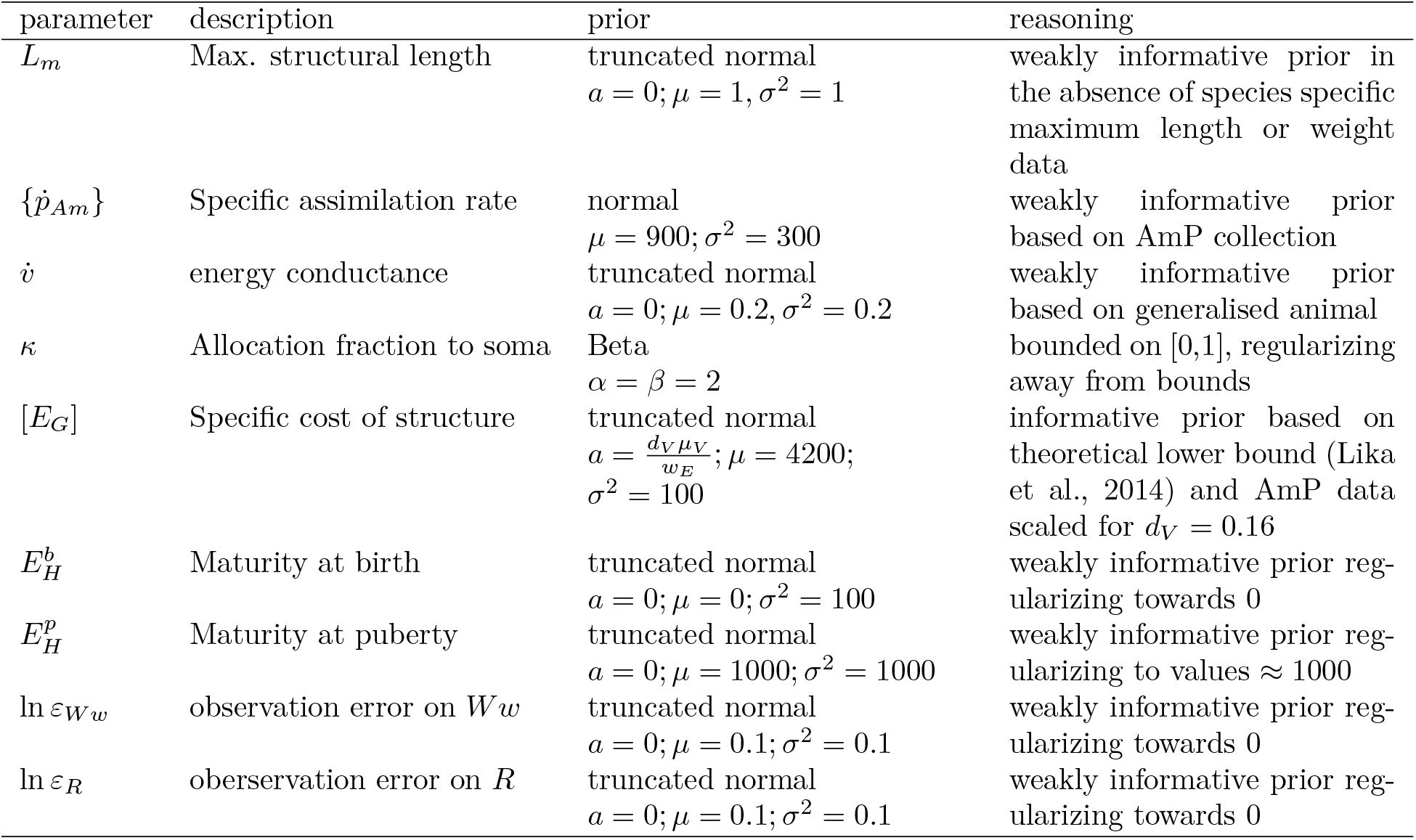
Parameters of the standard DEB model that were estimated in the earthworm case study, and their associated prior distributions and hyper-parameters.

**Table 3:**
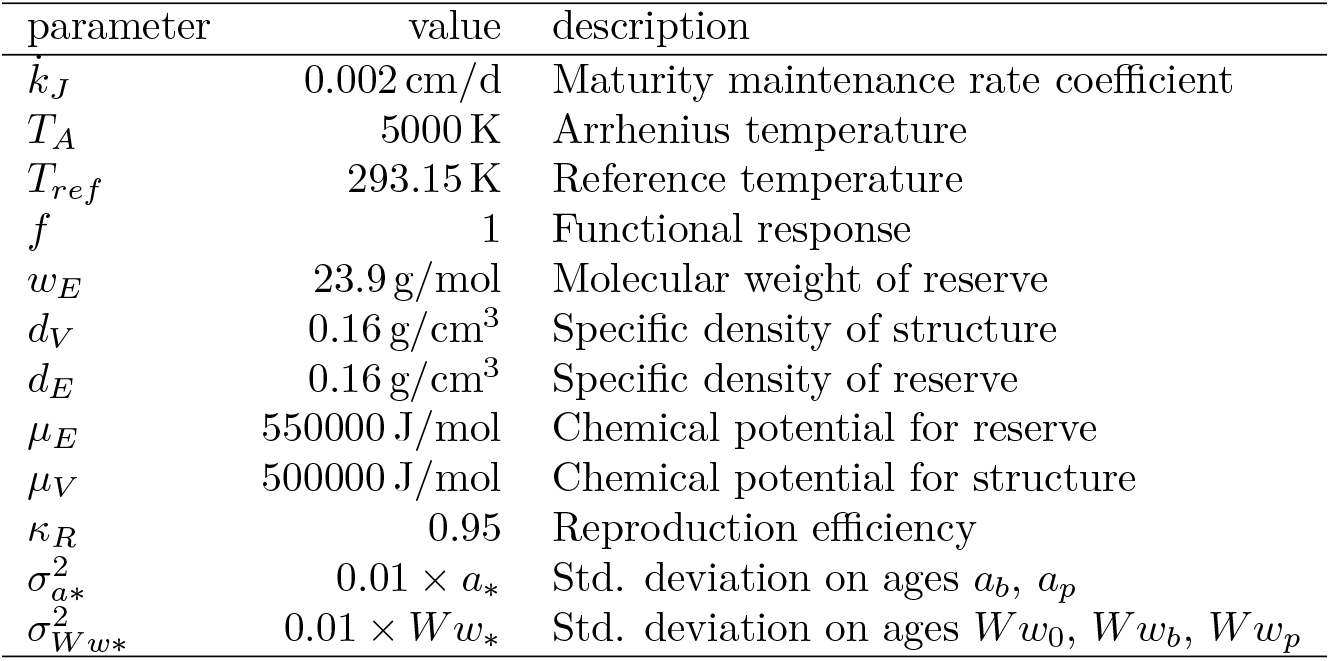
Parameters of the standard DEB model that were treated as fixed in the estimation procedure. Values are based on the AmP entry for *Lumbricus terrestris.* No variance estimates for ages *a_b_,a_p_* and weights *W_w0_,W_wb_,W_wp_* at stage transitions in *L. terrestris* were available in the literature. We therefore assumed an arbitrary fixed standard deviation 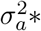 and 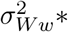 of 1% of the reported mean for those values. Additional parameters are defined in Appendix A.

For the sake of simplicity we did not estimate ageing parameters. DEB model parameters estimated from observations of earthworm growth (Butt, 1993) with DEBtool_M (https://github.com/add-my-pet/DEBtool_M) were obtained from the add-my-pet database (Marques et al., 2018). Rather than re-estimating the DEB parameters from the original data we simulated observations for this case study, by solving the DEB equations with the AmP parameters for this species, and adding random log-normal noise to the simulated trajectories. This allowed us to conduct inference on a set of known reference parameters for both the deterministic model, and the observation model, and thus to objectively assess parameter identifiability and the precision of the posterior parameter estimates independent of the inferential performance of DEBtool. The code for the simulation procedure is provided in the supplementary materials.

### 4.1 Bayesian parameter estimation

Treating the initial value for the scaled reserve density 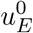 as parameter dependent (Kooijman, 2009) necessitates a recalculation of two of the initial values for the DEB model equations, the scaled length *l_b_* and maturity at birth 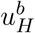, respectively, within the MCMC inference procedure. This computational step is currently only implemented in a development branch of `deBInfer`, which is provided in the supplementary materials.

Initial inference runs highlighted parameter identifiability issues, in particular a strong, nonlinear correlation between the maximum structural length *L_m_*, the specific assimilation rate 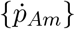 and the energy conductance *v* made it impossible to estimate the specific cost of structure [*E_G_*]. We tried to resolve this by using informative priors based on the empirical distribution of parameters in the AmP database (see below).

The strong parameter correlations further resulted in slow mixing of the MCMC chain, necessitating a large number of samples to explore the posterior distribution and achieve approximate convergence, as indicated by potential scale reduction factors < 1.1 (Brooks and Gelman, 1998). We therefore conducted inference for this model using a compiled ODE model implemented in C to make inference feasible in an acceptable amount of wall time. We ran three separate MCMC chains of 150000 samples each. We discarded the first 10000 samples of each chain as ‘burnin’ and thinned the remaining samples retaining every 10th sample.

#### 4.1.1 Prior distributions

Priors on the parameters were chosen to be weakly informative, based on the principle that priors should contain enough information to rule out unreasonable parameter values but not values that might make sense. Hard bounds were used only where dictated by DEB theory. We further aimed to translate the pseudodata approach of the covariation method (Lika et al., 2011) into our choices of prior distributions and their location and scale parameters. Specific prior choices are detailed in Table 2. Informative priors for 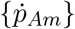 and [*E_G_*] were based on exisiting DEB parameter estimates. To this end we extracted DEB parameter values across taxa for all members of the kingdom Animalia represented in the AmP database using the prtStat function of AmPtool (Marques et al., 2018), and based informative priors on the interspecific means and standard deviations.

#### 4.1.2 Data model and likelihood

The state variables of the DEB model are abstract quantities that are not directly observable, but can be mapped to observable quantities using auxiliary parameters. We used the following mappings between the so-called zero-variate observable quantities and model states and parameters (see Tables 2 and 3 for definitions):

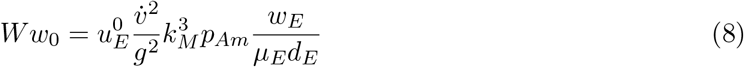

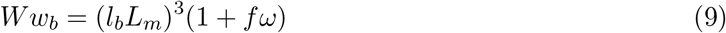

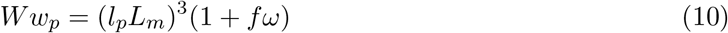

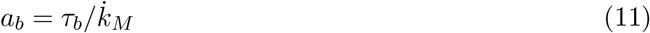

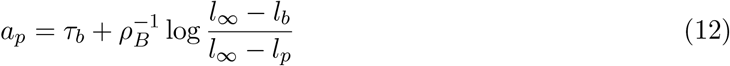

where 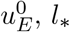, and *τ_b_* were calculated following Kooijman (2009).

Further, the time series of wet weight *Ww*(*t*) and reproductive output *R*(*t*) were mapped from the model using the equations

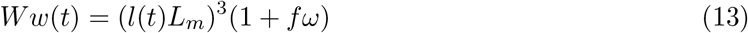

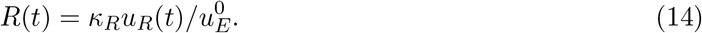

As in the first case study we assume multiplicative log-normally distributed error terms on the time-series observations of lengths and cumulative egg production, but truncated normal likelihoods on the zero-variate observations of weights and ages at the start of development, birth, and puberty.

This choice of likelihood offered a straightforward way to parameterize the likelihood, given that zero-variate data are often reported as means and standard deviations, while the truncation ensures that only positive values are allowable. The full likelihood of error at time *t* is therefore as follows

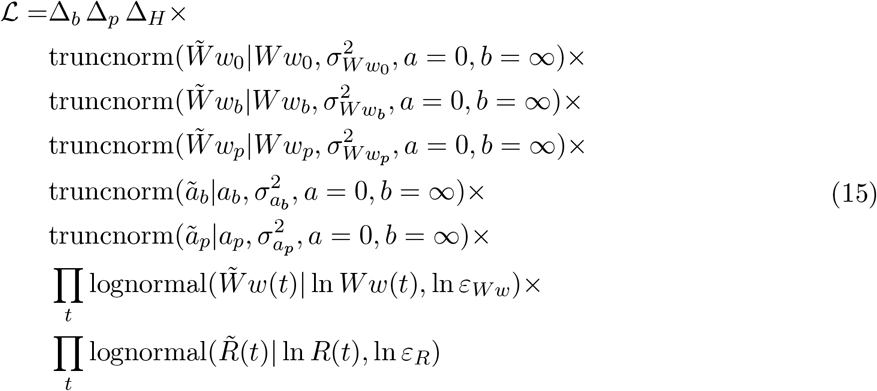

where further constraints Δ on the parameter space follow from theoretical considerations detailed in Lika et al. (2014) about the animal reaching the stage transitions at birth and puberty:

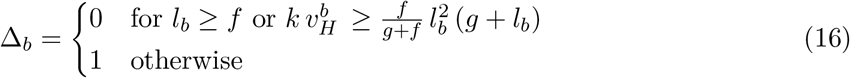

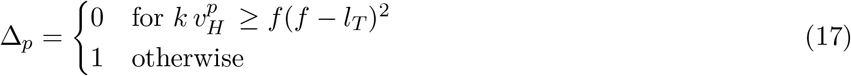

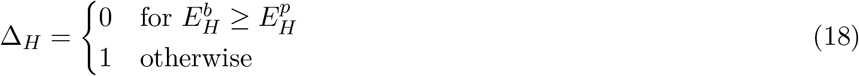

### 4.2 Results: Standard DEB Model

Initial inference runs highlighted parameter identifiability issues, in particular the strong, non-linear correlation between 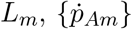 and 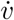 (see the banana-shaped contours in the joint density plots for the variables in Figure 5) made it impossible to estimate [*E_G_*], even when using informative priors based on the empirical distribution of parameters in the AmP database. We therefore present inferences for a model where [Eg] was fixed at the value of 4180 J/cm^3^ (Figure 6).

**Figure 5:**
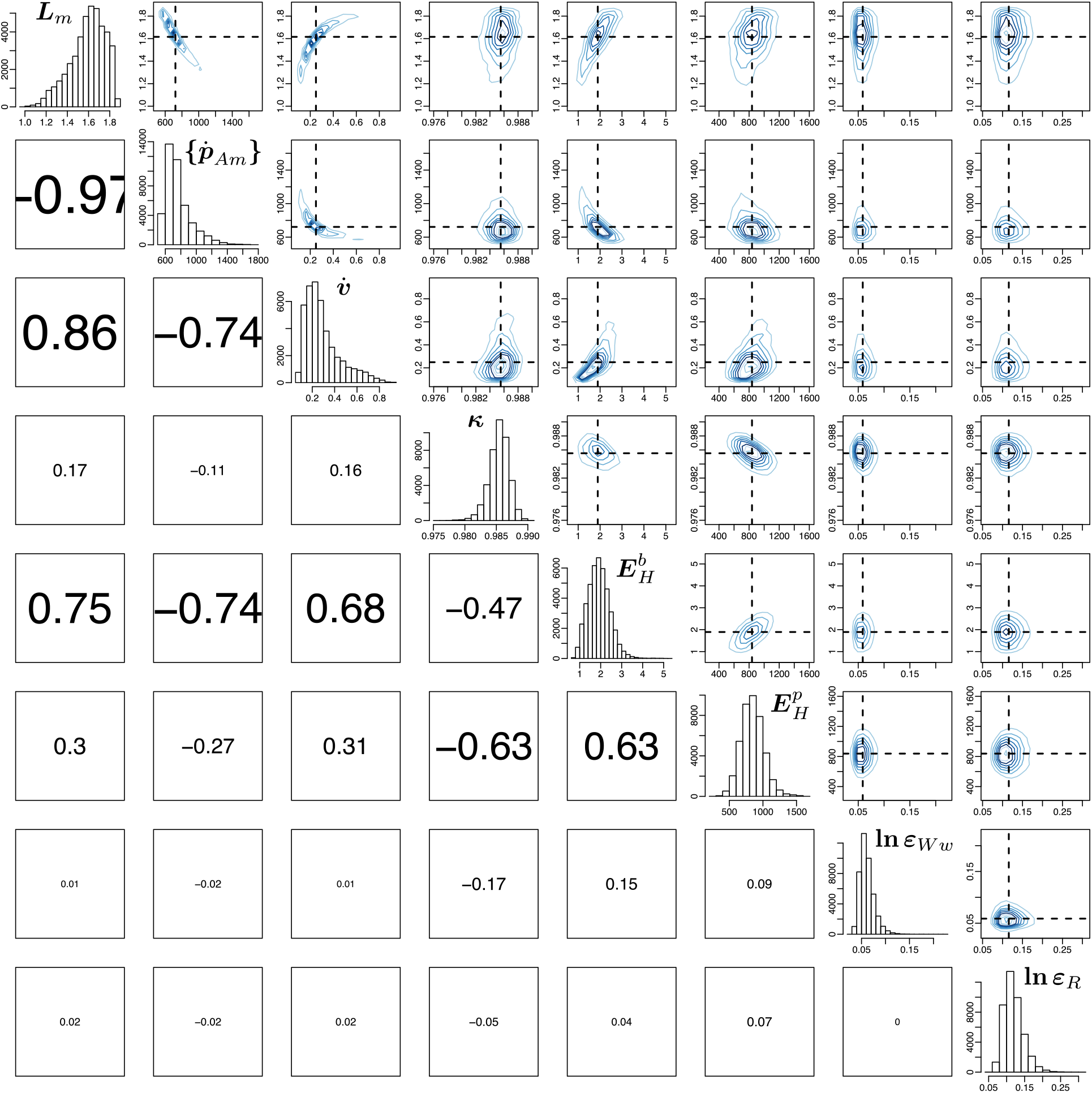
Pairwise correlations of posterior parameter estimates for the standard DEB model for the earthworm *Lumbricus terrestris.* The value for the specific cost of structure [*E_G_*] was fixed at 4180 J/cm^3^.

**Figure 6:**
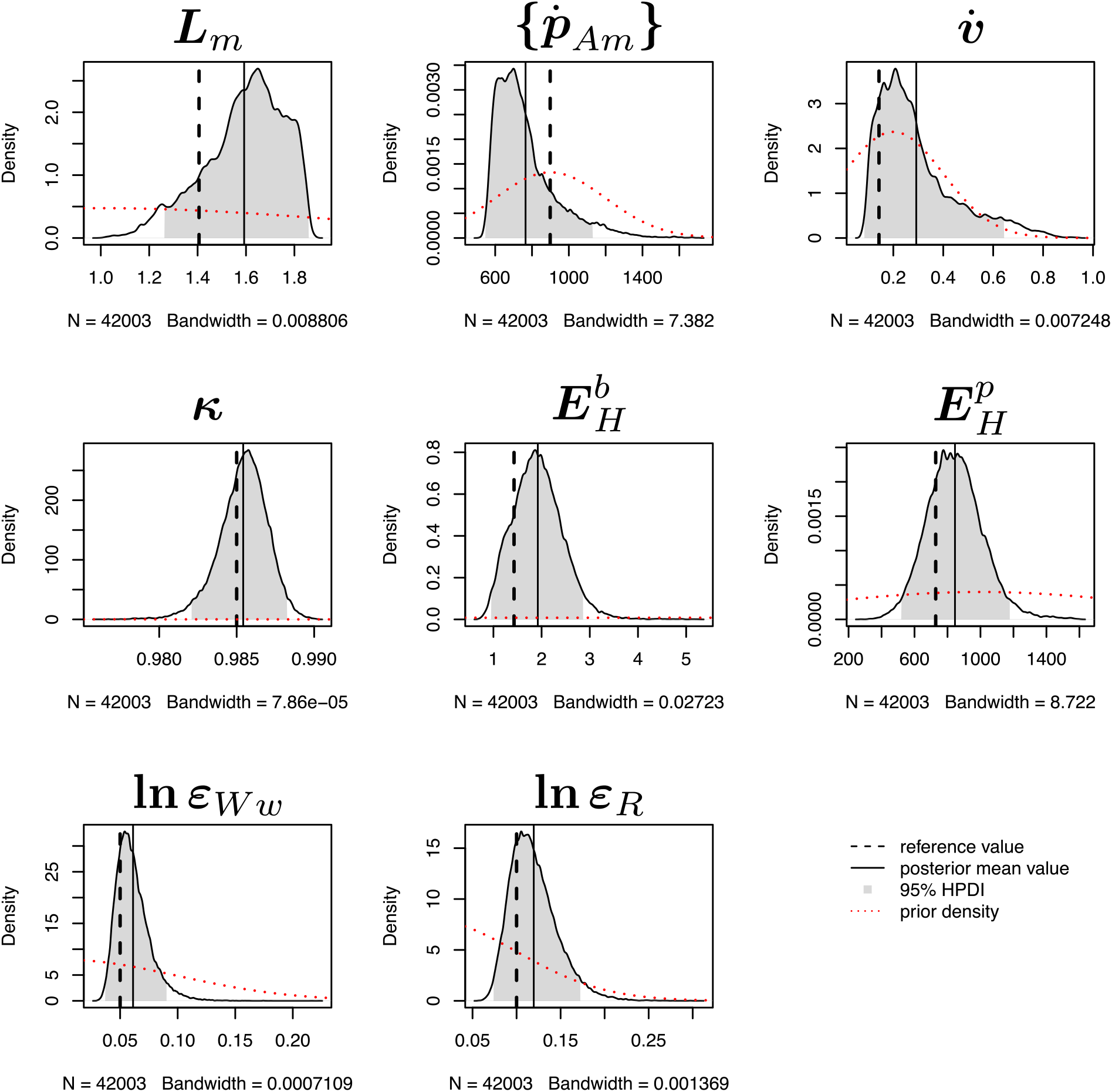
Even with a fixed value of the specific cost of structure [*E_G_*] the data likelihood provided little information about the values of energy conductance 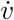 and the specific assimilation rate 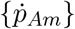, as is apparent from the substantial overlap between posterior and prior densities for these parameters. Despite this, the reference values of all free parameter were recovered within the 95% highest posterior density interval.

Despite the strong correlations, the AmP parameter values were recovered within the 95% highest posterior density interval, although the posterior means and modes diverged substantially from the AmP parameter values for 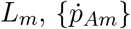, and 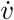, and to a lesser extent for 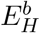 (Figure 6).

Posterior trajectories for the earthworm DEB model (Figure 7) further indicate that the parameter identifiability issues encountered here are likely a structural property of the model, rather than a result of poor statistical fit due to large observation errors. The posterior distribution of model trajectories is relatively narrow on the data scale, which is well constrained by the observations, but wide on the scale of the state variables.

**Figure 7:**
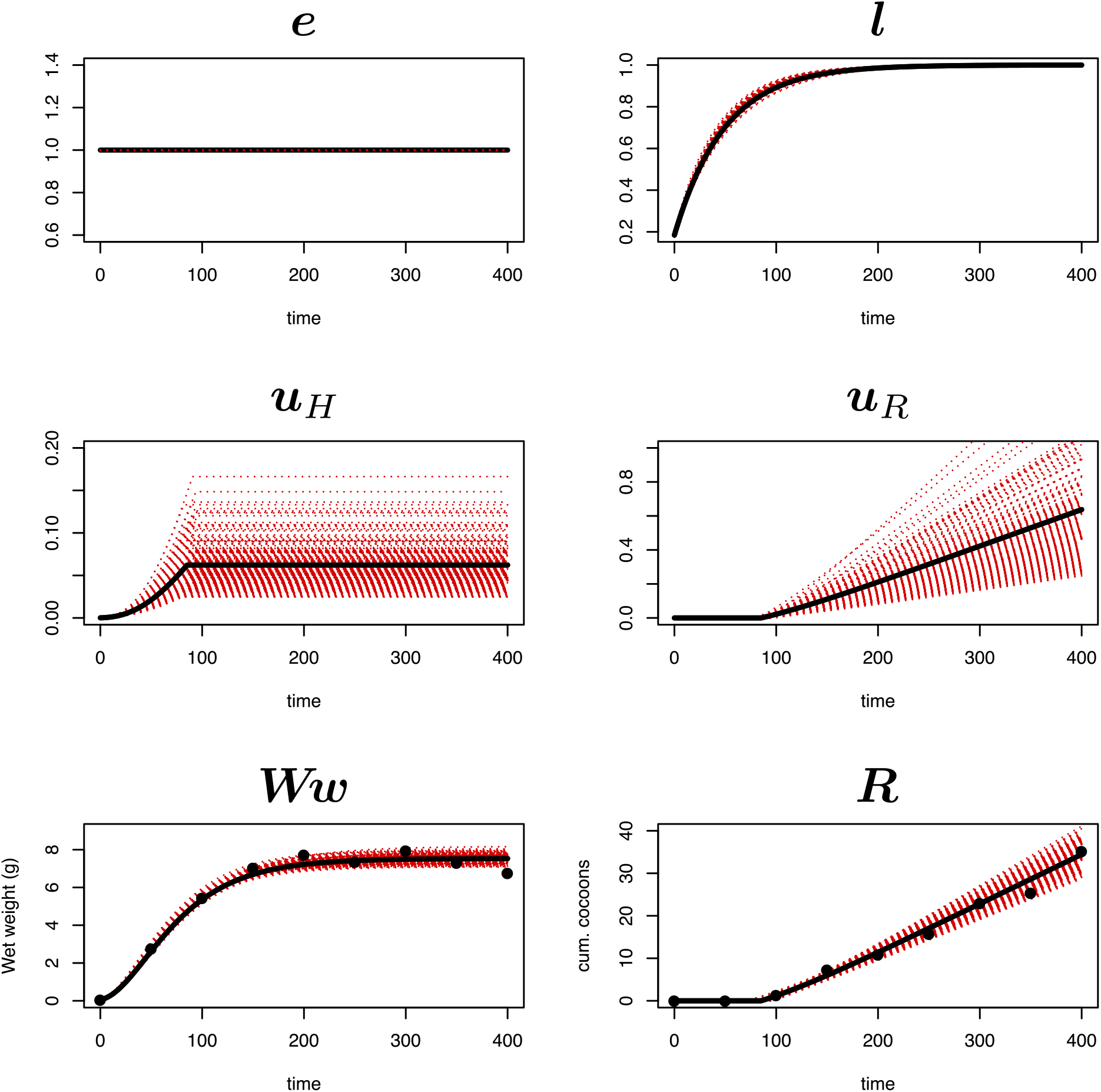
Posterior trajectories for the earthworm DEB model. The posterior distribution of model trajectories is relatively narrow on the data scale, but wide on the scale of the state variables. This indicates that the weak identifiability of several parameters is structural, rather than a consequence of poor statistical fit. The red dashed lines are a random sample of 500 posterior trajectories. The black solid line represents the simulated trajectories which were used to generate the noisy observations (black dots) used as data in the inference procedure.

## 5 Discussion

DEB theory and related bioenergetic models such as the DEBkiss framework have the potential to unify biological theory across levels of organization (Nisbet et al. 2000). However, to realize this potential, robust methods are needed to connect the theoretical predictions with the inherently variable and noisy biological data.

We here demonstrate how DEB and DEBkiss parameters can be estimated in a Bayesian framework, a coherent, effective, and well-established approach, using the freely available deBInfer package in R. The Bayesian approach is particularly useful for DEB models as it provides a fairly straightforward way to incorporate prior information and at the same time provides a means to quantify uncertainty in parameters and predictions. DEB theory in its very core is focused on the individual animal, and the role of individual trait heterogeneity is increasingly recognized as an important factor underlying ecological dynamics (e.g. Cam et al., 2002; Vindenes et al., 2008; Jenouvrier et al., 2015). The problem of inter-individual variation and thus dependence in observations remains an important source of bias for inference in DEB models. The Bayesian approach, in principle, provides a conceptually straightforward avenue for hierarchical inference, thereby opening a door to better understanding causes and effects of individual heterogeneity of metabolic traits. However, the computational implementation of such a hierarchical inference is challenging, and currently exceeds the capabilities of the `deBInfer` software. Progress towards more general tools for this purpose is being made (Carpenter et al., 2017; Krauss and Schuppert, 2017), and should provide great opportunities to further develop inferential approaches for DEB models. Our results furthermore highlight the difficulty of identifying DEB model parameters when taking into account measurement uncertainty, which serves as a reminder that fitting these models requires statistical caution (Billoir et al., 2008a). In particular, we were not able to simultaneously estimate the same number of parameters for the standard DEB model for *Lumbricus terrestris* as are presented in the corresponding AmP entry, even when using priors based on AmP information. Further work should employ additional datasets to further elucidate the influence of data availability (as in the number of measured traits) and quality (as in the precision of observations) on practical parameter identifiability in the standard DEB model when assuming observation error.

Any statistical inference procedure involves arbitrary choices made by the modeller, and both the Bayesian approach presented here, and the DEBtool procedure make use of distributional assumptions, as well as expert opinion to constrain the parameter estimation. The former through the choice of particular prior distributions and likelihoods, the latter by using pseudodata and setting weight coefficients for the least-squares estimation. Furthermore, the weighted least-squares method underlying the DEBtool estimation procedure does in principle provide variances and approximate covariances on parameter estimates, however, these are rarely if ever reported, and are not currently part of the AmP database. To better understand the uncertainties of parameter estimates we would encourage all DEB practitioners to report choices made to constrain the parameter estimation, as well as variance and covariance estimates for estimated parameters.

## 6 Acknowledgements

The authors were supported by the US National Science Foundation (Grant PLR-1341649). We thank Sadie Ryan, the participants of the 5th International Symposium on Dynamic Energy Budget Theory, and two anonymous reviewers for comments on earlier versions of this work.

## 7 Code and Data availability

The earthworm data and parameters were obtained from the add-my-pet library (AmP; http://bio.vu.nl/thb/deb/d entry *Lumbricus terrestris* version 2015/12/07,

Code and derived data sets for this paper are available online.

- DEButilities 0.1.0 is archived on zenodo (https://doi.org/10.5281/zenodo.1162331)
- deBInfer 0.4.2 is available on CRAN (https://CRAN.R-project.org/package=deBInfer)
- deBInfer 0.4.1.9000-recalc-inits is available on github (https://github.com/pboesu/debinfer/tree/recalc-inits)
- Simulation and inference code for this paper are archived on zenodo (http://doi.org/10.5281/zenodo.1298407)

## A The standard DEB model

Dynamics for the scaled standard DEB model for *e* > *l* > *l_b_* following Table 2.5 in Kooijman (2010).

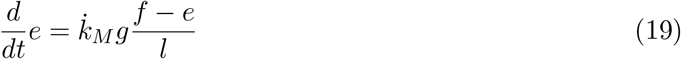

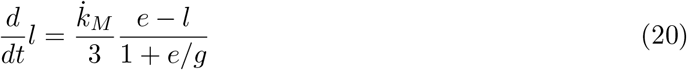

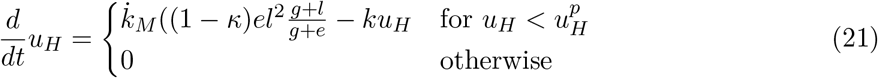

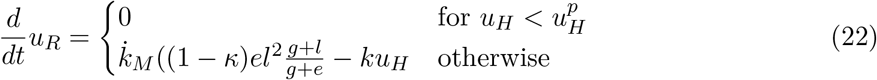

**Table 4:**
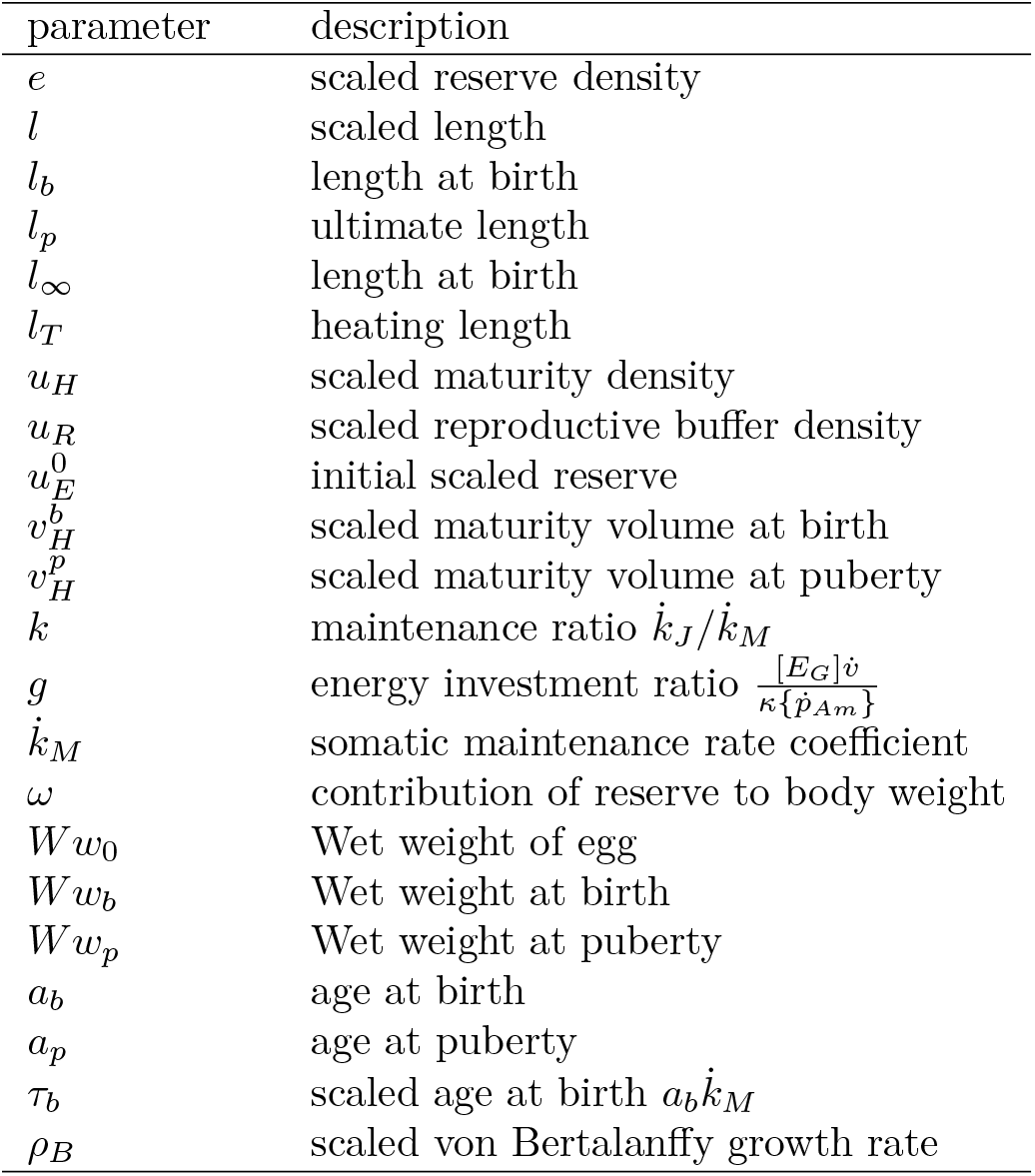
Definitions of state variables and parameters of the standard DEB model not detailed in Tables 2 or 3.

